# Spatiotemporal analysis of autism gene enrichment implicates cortex, thalamus, and hypothalamus

**DOI:** 10.64898/2026.05.14.724487

**Authors:** Xuran Wang, Yue Li, David M. Young, Alicia Ljungdahl, Chimmi Dema, Narjes Rohani, Tomas Nowakowski, Kathryn Roeder, Stephan J. Sanders

**Affiliations:** Seaver Autism Center for Research and Treatment, Icahn School of Medicine at Mount Sinai, New York, NY 10029, USA; Department of Psychiatry, Icahn School of Medicine at Mount Sinai, New York, NY 10029, USA; Department of Genetics and Genomic Sciences, Icahn School of Medicine at Mount Sinai, New York, NY 10029, USA; Department of Statistics and Data Science, Carnegie Mellon University, Pittsburgh, PA 15213, USA; Department of Psychiatry and Behavioural Sciences, University of California, San Francisco, San Francisco, CA 94158, USA; Department of Paediatrics, Loma Linda University, Loma Linda, California, USA 92354; Institute of Molecular and Cell Biology, A*STAR, Singapore 138673, Singapore; Institute of Developmental and Regenerative Medicine, Department of Paediatrics, University of Oxford, Oxford, OX3 7TY, UK; Department of Neurological Surgery, University of California San Francisco, San Francisco, CA 94143, USA; Weill Institute for Neurosciences, University of California San Francisco, San Francisco, CA 94158, USA; Department of Anatomy, University of California San Francisco, San Francisco, CA 94143, USA; Eli and Edythe Broad Centre of Regeneration Medicine and Stem Cell Research, University of California San Francisco, San Francisco, CA 94143, USA; Department of Computational Biology, Carnegie Mellon University, Pittsburgh, PA 15213, USA; New York Genome Centre, New York, NY 10013, USA

## Abstract

Autism spectrum disorder (ASD) is a highly heritable neurodevelopmental disorder. Sequencing analyses have identified 185 ASD-associated genes, which implicate neurons, but the specific brain regions through which these neurons influence neurodevelopment remain unclear. Here, we integrate over one million single-cell RNA sequencing profiles from 20 regions of the developing human brain (4–23 post-conceptual weeks) using a new framework, STARMAPS (Sparse Task-specific Analysis for Revealing Molecular Associations in Particular Single-cell datasets). STARMAPS accounts for coordinated regional and developmental perturbations in gene expression, enabling robust cross-region comparison. We replicate prior findings that ASD-associated genes are enriched in excitatory neurons during mid-fetal development, and we extend these results to reveal distinct spatial signatures. Across 26 excitatory neuron subtypes, six clusters showed significant enrichment for ASD-associated genes. These clusters localize to both cortical and subcortical regions, including the motor, temporal, and visual cortex, as well as the thalamus and hypothalamus. Our findings support a major role for excitatory neurons across distributed brain circuits, implicating previously underappreciated subcortical structures in ASD etiology. By providing a statistically rigorous framework for spatiotemporal integration of single-cell data, STARMAPS enables refined mapping of molecular vulnerability across the developing human brain.

## Introduction

Autism spectrum disorder (ASD) is a highly heritable condition diagnosed in 3.22% of children.[1, 2] Exome sequencing of large cohorts has identified numerous genes associated with ASD (185 with false-discovery rate [FDR] ≤ 0.05).[3] Individuals with rare variants in these genes often have substantial impairments and severe co-morbidities, including global developmental delay and early-onset seizures. Many of the ASD-associated genes play a role in transcriptional regulation or neuronal communication.[4] These rare variants act on a polygenic genetic background that also contributes to phenotypes;[5, 6] a few common genetic variants mediating this polygenic background have been identified and appear to target convergent sets of genes and biological pathways as those containing rare variants.[7, 8] The etiology of ASD remains unknown, however, the causal relationship of the genes identified provides an entree to exploring when, where, and how these variants influence human behaviors.

Some brain disorders are associated with specific brain regions, for example, degeneration of dopamine neurons in the substantia nigra of the basal ganglia causes Parkinson’s Disease.[9] The brain regions that underlie ASD remain unknown and is a longstanding, critical challenge.[10] Resolving this question is critical to designing *in vivo* experiments to understanding the role of specific genes in these disorders, addressing circuit-based experiments, and assessing the biodistribution and dosage of novel therapeutics or targeted neuromodulation. The brain regions underlying numerous neurological traits have been resolved through the identification of individuals with similar symptoms due to localized brain lesions (e.g., congenital malformations, tumors, cerebrovascular accidents). While such lesions are observed more frequently in individuals with ASD in general, they do not pinpoint a consistent locus.[11–13] Similarly, neuroimaging approaches, including structural and functional data, suggest subtle differences between the brains of individuals with ASD and typically developing controls, but without a clear consensus on the brain regions identified.[14–16]

Single-cell RNA sequencing (scRNA-seq) data provides an opportunity to assess whether specific cell types express ASD genes to a higher degree, following the logic that a gene must be expressed to impact cell function. Applying these gene enrichment analyses to the developing human cortex have strongly implicated excitatory and, to a lesser extent, inhibitory neurons.[3, 8] Bulk tissue RNA-seq datasets from the developing *postmortem* human cortex implicate the second and third trimester of fetal development, though this may also reflect the relatively high proportion of neurons at these stages.[3, 8] ASD-associated genes are also enriched among dysregulated genes in the *postmortem* cortex of individuals with ASD.[17–19]

scRNA-seq data from multiple regions of the developing human brain provide the opportunity to assess which regions are enriched for ASD-associated genes and, therefore, are more likely to be involved in ASD etiology.[20, 21] However, we expect profound spatiotemporal differences within and between cell types and this creates a challenge for data integration in these diverse data. In essence, the task is akin to overcoming a batch effect; however, it is more subtle, since a regional or temporal effect involves a global and simultaneous perturbation of many genes. To address these challenges, we developed the Sparse Task-specific Analysis for Revealing Molecular Associations in Particular Single-cell datasets (STARMAPS) framework, which builds on the previously described cFIT framework, but it brings greater flexibility and interpretability.[22] STARMAPS uses the concept of task-specific subspace perturbation, which has been utilized in language model and computer vision applications. Overall, this model also provides an intuitive approach that allows for seamless inference about gene expression within cell types, while controlling for the regional effect.

Applying STARMAPS to scRNA-seq data collected from over 20 dissected regions of the developing human brain between 4 and 23 post-conceptual weeks (PCW) [20, 21] and we replicate the enrichment of ASD-associated genes in neurons and mid-fetal development. Controlling for regional effects, we identify six ASD-enriched clusters of excitatory neurons. Considering the original locations of the neurons within these six clusters, we see involvement of both cortical and subcortical regions, including the thalamus and hypothalamus.

## Results

### Single-cell expression across the developing human brain

Two recently published datasets[20, 21] enable spatiotemporal assessment of ASD-associated gene enrichment (Fig. 1). Eze *et al*. details single-cell RNA-seq data generated with the 10X Genomics Chromium platform from ten individual brains in the first trimester (Carnegie stages [CS] 12 to 22, corresponding to 4-7 PCW).[20] Approximately 289,000 high-quality cells were collected from 17 anatomically distinct dissected brain regions (Fig. 1, Supplementary table 1). Bhaduri *et al*. also describes 10X Genomics Chromium single-cell RNA-seq data but from eleven individual brains in the second trimester (12-23 PCW). The ∼700,000 high-quality cells were obtained from 17 anatomically distinct dissected brain regions (Fig. 1). Together, these two datasets capture about a million cells across key structures in the forebrain, midbrain, and hindbrain throughout the first and second trimester (4 to 23 PCW).

**Figure 1.**
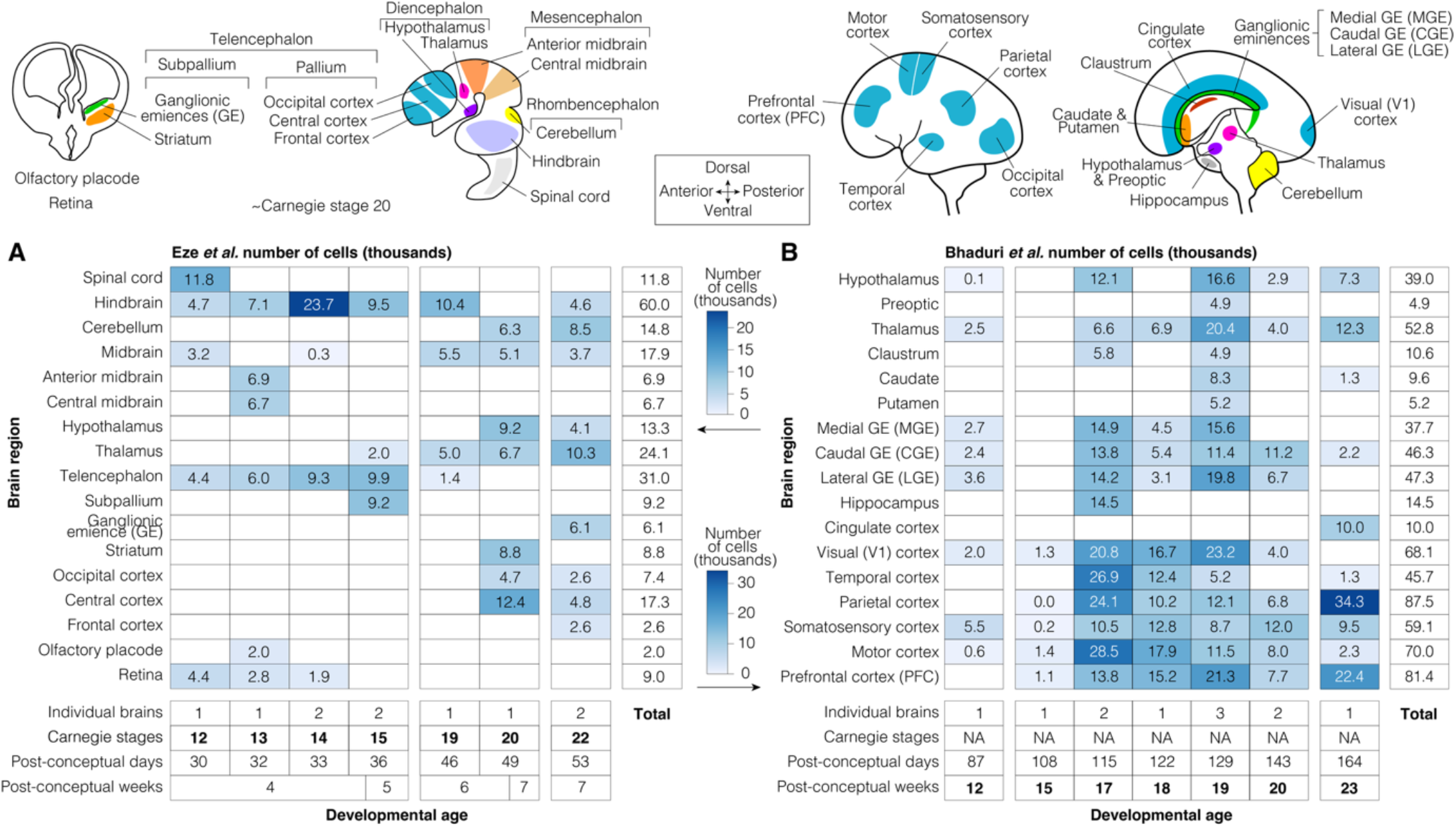
Overview of samples and cells. **A)** About 289,000 high-quality cells were collected from 17 brain regions across ten individual brains in Eze *et al*.[20]. The distribution of cell counts is shown by developmental stage (x-axis) and brain region (y-axis); darker shades represent more cells. **B)** Equivalent data from ∼700,000 high-quality cells from 17 brain regions across eleven individual brains from Bhaduri *et al*.

### ASD-associated gene enrichment implicates mid-fetal development

Our first goal is to assess the relative enrichment of ASD-associated gene expression across stages of prenatal brain development. Due to gaps in sample availability across developmental trajectories (e.g., CS15-19, 7-12 PCW, Fig. 1) and the dramatic changes in gene expression across these developmental stages[23], we elected to assess the dataset in three groups: CS12-15, CS19-22, and 12-23 PCW.

Within each of the three developmental groups we assessed ASD enrichment, i.e. the extent to which 255 ASD-associated genes (FDR ≤ 0.1) were expressed, following the logic that higher relative expression represents a higher probability of contributing to ASD neurobiology. Enrichment of ASD genes was assessed on a per cell basis by calculating the odds ratio and p-value (Fisher’s Exact Test) between expressed and non-expressed genes for ASD and non-ASD protein-coding genes (Supplemental Methods). The percentage of cells that are enriched (OR > 1, P≤0.05) for ASD-associated genes is shown in Fig. 2 for each of the three developmental groups, split by brain region.

**Figure 2.**
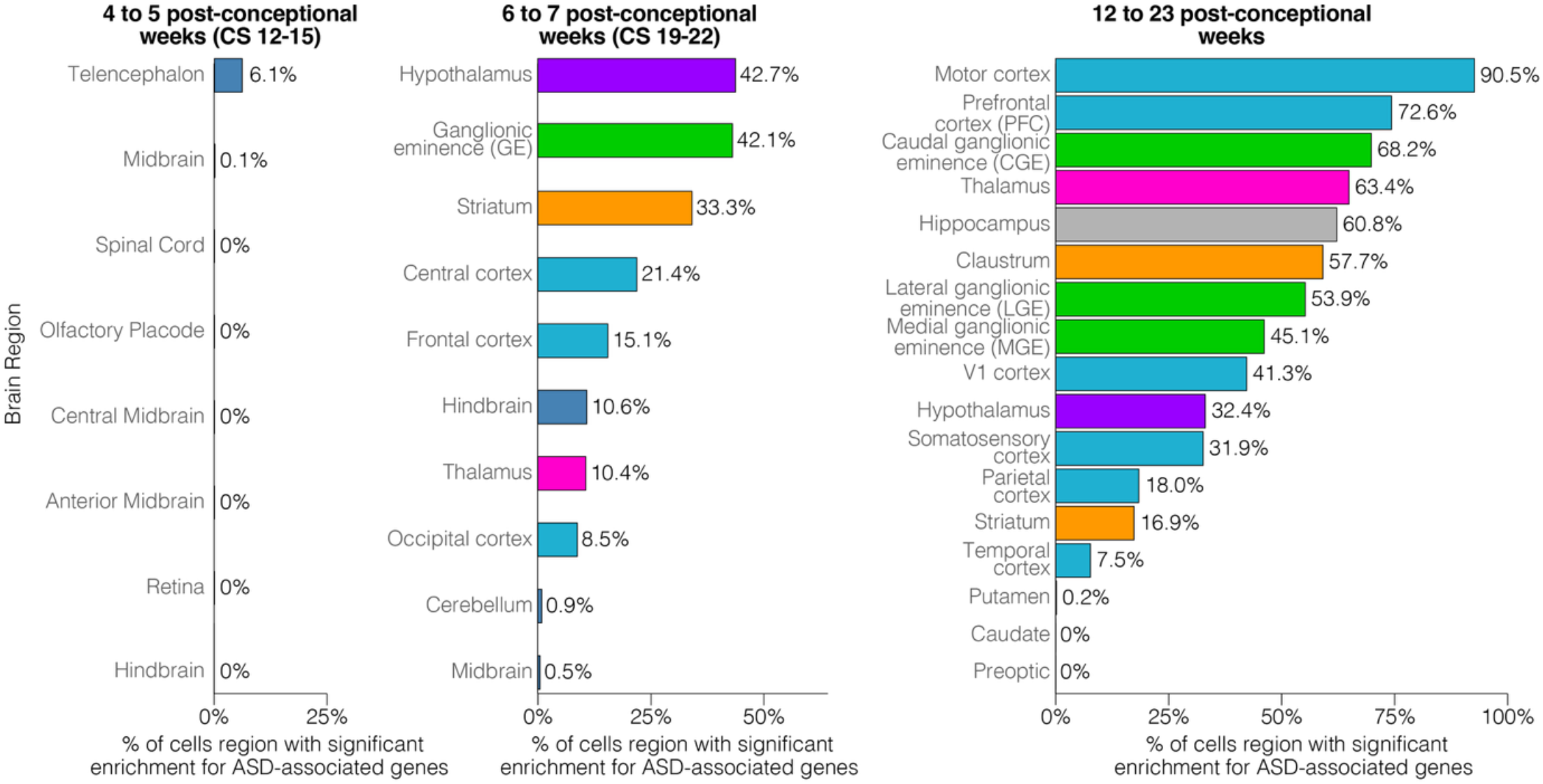
Enrichment of ASD-associated genes across prenatal development. Enrichment of ASD-associated genes was assessed on a per cell basis (Fisher’s Exact Test) and the percentage of ASD enriched cells (x-axis) is show as a percentage in each dissected brain region (y-axis) for the three developmental stages (left to right).

The earliest developmental group (CS12-15, 4-5 PCW) has few ASD-enriched cells (Fig. 2); those that are observed are restricted to the telencephalon (6.1%) and midbrain (0.1%). By 6-7 PCW (CS19-22), substantially more cells are enriched for ASD-associated genes. Subcortical regions show the greatest degree of enrichment, hypothalamus (42.7%), ganglionic eminence (42.1%), and striatum (33.3%), followed by frontal (21.4%) and central (15.1%) cortex. However, by 12-23 PCW, ASD enrichment is dramatically increased, especially in the cortex, with 90.5% of motor cortex cells being enriched for ASD genes followed by 72.6% of prefrontal cortex cells. Subcortical regions still have many ASD-enriched cells at this later developmental stage, including the caudal ganglionic eminence (68.2%), thalamus (63.4%) and hippocampus (60.8%). Thus, while ASD genes begin to be enriched in cells in the 1^st^ trimester, the enrichment is much more substantial in the 2^nd^ trimester. This result is in keeping with prior analyses of bulk and single-cell transcriptomic data.[3, 8, 24] We therefore focused our further analysis on this later ASD gene-enriched developmental stage (12-23 PCW) that represented over 70% of the cells in this initial assessment.

### ASD gene enrichment across cell types and regions in the 2^nd^trimester brain

Within the 2^nd^ trimester data, gene expression varies by brain region, developmental stage, sample, and cell type, creating a challenge for data integration. Correcting for these effects needs to account for a global and coordinated perturbation across many genes, rather than considering each gene independently. We therefore developed STARMAPS, a more flexible and interpretable extension of the cFIT framework.[22] STARMAPS uses the concept of task-specific subspace perturbation, which has been utilized in language model and computer vision applications.[25–27] Building upon the non-negative matrix factorization framework, we assume that each region or temporal window is associated with a latent factor space. We then regularize directly on the (approximate) subspace distance between those factor spaces. STARMAPS provides more freedom on parameters, while facilitating control of task-specific perturbation through modification of the regularization parameters. Overall, this model also provides an intuitive approach to assess the degree of perturbation for each region/temporal, compared to the shared latent space, and the distances between latent spaces for regions (Fig. S1C, S1D).

To define cell types and brain regions enriched for 185 ASD genes, we performed gene-set enrichment analysis within these cell-region subgroups, using cell labels from the original description.[20, 21] We used STARMAPS to correct for batch effects and development changes, but to preserve regional and cell type differences (Fig. 3A/B). In keeping with prior analyses,[3, 8, 24] we observed substantial and significant enrichment for excitatory neurons across numerous brain regions (Fig. 3C); all brain regions considered showed a positive degree of ASD gene-enrichment within excitatory neurons, though not all reached the significance threshold (Fig. 3C). Inhibitory neurons showed a positive degree of ASD gene-enrichment in most regions, with parietal cortex, claustrum, and preoptic regions reaching significance. Some cortical regions were also enriched for ASD-associated gene in intermediate progenitor cells (IPC) (Fig. 3C).

**Figure 3.**
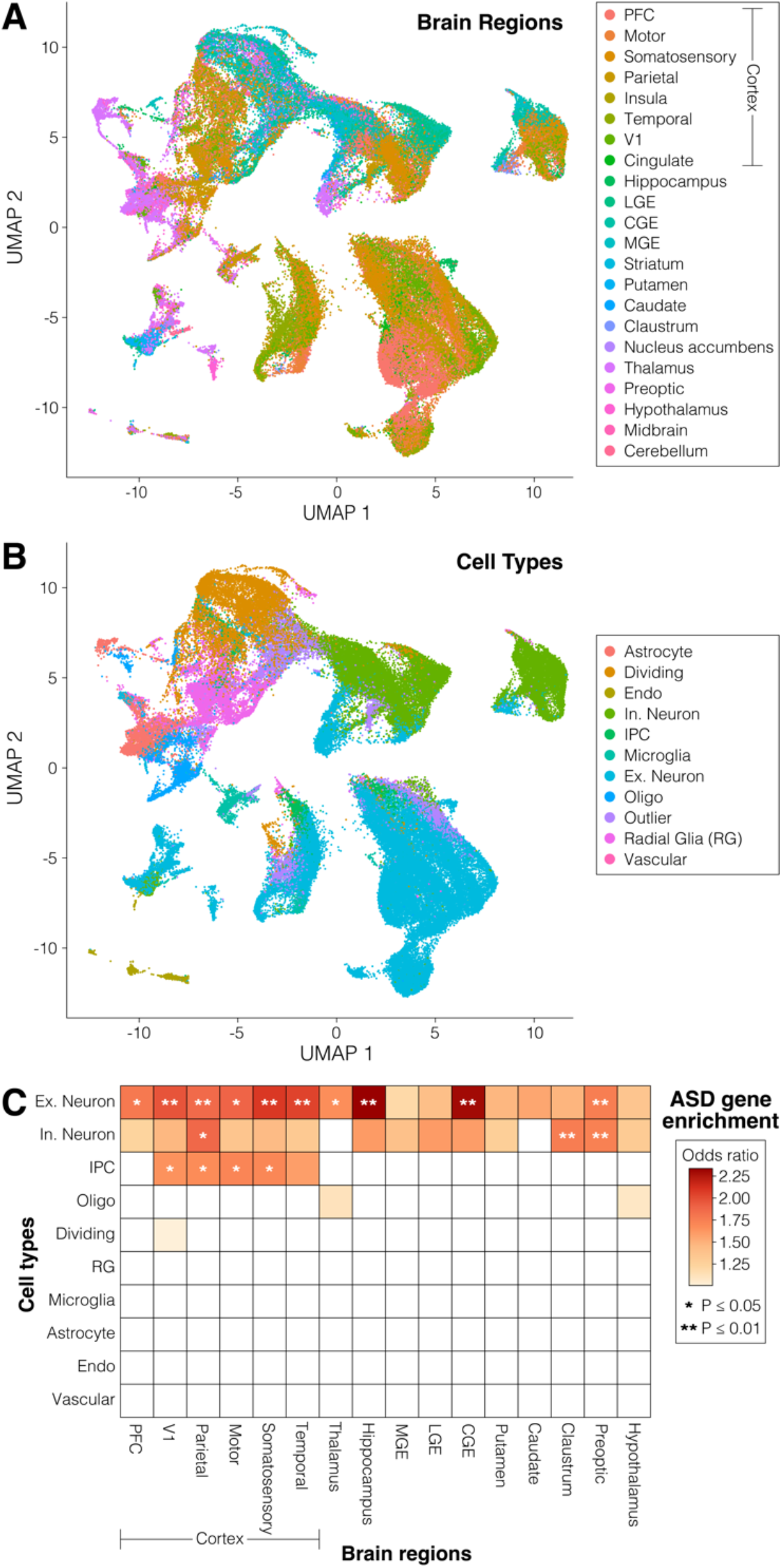
Integrated 2^nd^ trimester single-cell data. **A)** A UMAP plot is generated from the single-cell data, processed by the STARMAPS framework. Cells are colored by dissected brain region. **B)** The same UMAP, colored by cell-type label from the initial description of the dataset.[21] **C)** A heatmap showing the degree of enrichment for 185 ASD-associated genes in the cells from Panels A and B, in data corrected for sample, batch, and developmental stage. Shade represents the magnitude of positive odds ratios; stars represent the P-value.

### ASD gene enrichment across 2^nd^ trimester excitatory neurons

Given the strong enrichment for ASD-associated genes in 2^nd^ trimester excitatory neurons (Fig. 3C), we focused further analyses on this subset of the data. Our objective was to identify if the enrichment was driven by a specific subset of excitatory neurons and, if so, the nature of these excitatory neurons. Since there are numerous types of excitatory neuron, we used STARMAPS to generate a matrix across all excitatory neurons, correcting for sample/batch/developmental stage and for brain region.

Within the STARMAPS-corrected excitatory neurons, we used the Seurat nearest neighbor clustering algorithm[28] to identify excitatory neuronal subclasses. An appropriate number of neuronal subclusters (26) was determined by applying MRTree,[29] which generates a tree structure across multiple resolution levels to assess cross-level cluster stability (Fig. S2).

We next assessed the degree to which the 26 excitatory neuronal subclusters were enriched for 185 ASD-associated genes. All 26 subclusters showed a trend towards such enrichment (Fig. 4A), in keeping with the result at the cell type/regional level (Fig. 3C). However, six subclusters stood out for being markedly and significantly enriched amongst the excitatory neurons (P_adj_ ≤ 0.05, Fisher’s Exact Test, Bonferroni correction, Fig. 4A). We therefore sought to understand the characteristics of the excitatory neurons that composed these subclusters.

**Figure 4.**
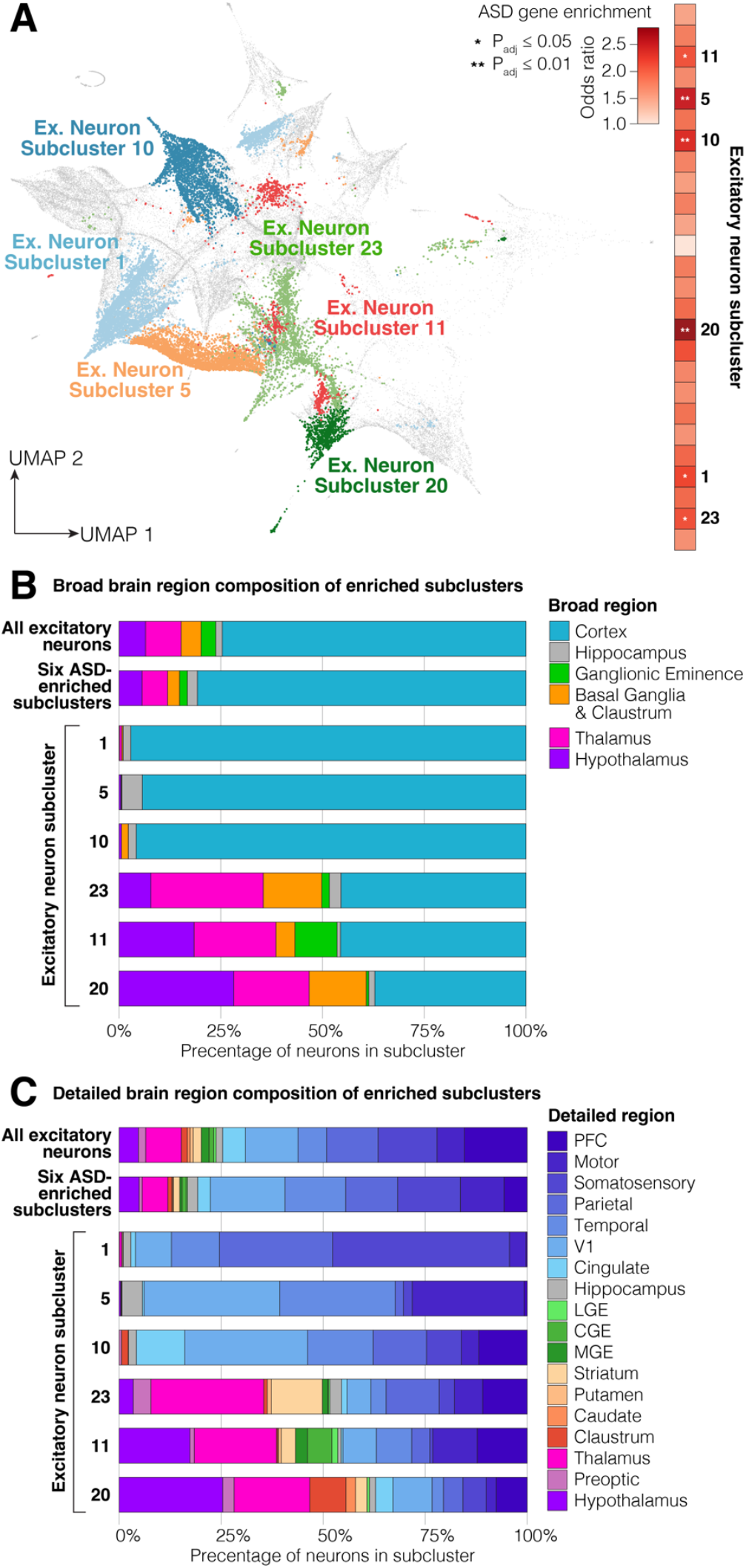
ASD-enriched excitatory neuronal subclusters. **A)** UMAP of all excitatory neurons identified 26 subclusters from the 2^nd^ trimester data, six of which are enriched for 185 ASD genes, as shown on the heatmap to the right. **B)** Brain regions of the original dissections for the six ASD-enriched excitatory neuronal subclusters and comparison to all excitatory neurons. C) The data in panel B are shown with more detailed brain region subsets. Statistical tests: A: Fisher’s Exact Test with Bonferroni correction.

A clear distinction was evident when considering the brain regions that the neurons were dissected from. Three subclusters (1, 5, and 10) were composed almost exclusively from cells dissected from cortical regions or hippocampus (Fig. 4B), whilst in the other three subclusters (11, 20, and 23) neurons from the thalamus and hypothalamus were substantially over-represented (Fig. 4B).

### ASD gene enrichment in 2^nd^ trimester cortical excitatory neurons

While substantial evidence supports the role of the cortex in ASD,[8, 30, 31] it is hard to distinguish whether this reflects greater availability of data due to the presumed role of the cortex or that the cortex genuinely plays more of a role than other brain regions. This multiregional analysis, ASD enrichment supports a major role for the cortex, with three of the ASD-enriched clusters composed almost exclusively from cells dissected from cortical regions (Subclusters 1, 5, and 10; Fig. 4B). To assess which cortical regions are involved, we annotated the cell composition of each subcluster by detailed dissected region (Fig. 4C, Fig. S3).

While the prefrontal cortex (PFC) was over-represented in subcluster 5, overall, a lower proportion of excitatory neurons from PFC were in the ASD-enriched subclusters than in all 26 subclusters (Fig. 4C). The cortical regions with the greatest over-representation were visual/V1 (Subclusters 5 and 10), temporal (Subcluster 5), and motor/M1 (Subcluster 1), and cingulate (Subcluster 10).

To assess cell type composition, we considered the marker genes in the untransformed data (Fig. S4). While detailed transcriptional atlases are available for cell types in adult human cortex, there are fewer prenatal resources. For example, 84-97% of the cells were ‘miscellaneous’ with the MapMyCell tool.

Subclusters 1 and 5 share expression of many of the marker genes (Fig. S4), despite being distinct in the hierarchical MRTree analysis, suggesting underlying broad transcriptional differences. These marker genes include *LHX2* and *NRP1* in clusters 1 and 5, *EMX2* in cluster 1, and *MEIS2* in cluster 5, suggesting these represent intratelencephalic (IT) neurons in the upper cortical layers (2/3). In contrast, marker genes in subcluster 10 include *FOXP1, SATB2*, and *RORB*, consistent with IT neurons in cortical layers 4/5.

Cortical samples are well represented across the developmental stages (Fig. 1), so we also considered the original developmental stages of the excitatory neurons making up these three ASD-enriched cortical clusters. We observed the greatest degree of ASD-associated gene enrichment at GW18 (81.0% of excitatory neurons ASD enriched) and GW19 (75.5%) compared with earlier stages (14.3% at GW14, 30.4% at GW17) or later stages (35.7% at GW20, 52.6% at GW22, 47.1% at GW25).

### ASD gene enrichment in 2^nd^ trimester subcortical excitatory neurons

The paucity of transcriptional data outside the human cortex has limited this line of inquiry for subcortical regions and, since many subcortical regions have lower neuronal density than the cortex, regional bulk tissue data could miss important signals. Therefore, it is notable that excitatory neurons across multiple subcortical regions are overrepresented in three of the ASD-enriched subclusters (11, 20, and 23, Fig. 4B, 4C, Fig. S3).

Subcluster 20 shows the greatest degree of enrichment for ASD-associated genes (Fig. 4A). The majority of neurons (64%) are subcortical, with over 25% of the total coming from the hypothalamus; overrepresentation is also seen in the thalamus and claustrum. Accordingly, we observe high expression of *SLC17A6* (VGLUT2), a marker of thalamic and hypothalamic regions.[32] The neurons dissected from hypothalamic regions are highly enriched for ASD genes, with 91.4% reaching nominal significance. To assess whether a specific nucleus of the hypothalamus contributed these neurons, we assessed the enrichment of subcluster marker genes in HypoMap, a unified single cell atlas of the murine hypothalamus.[33] We observed enrichment for genes from Subcluster 20 across multiple hypothalamic nuclei, especially the ventromedial hypothalamic nucleus, the median eminence, the dorsomedial nucleus of the hypothalamus, and the lateral preoptic area (Fig. S5). An equivalent reference set was not available for the thalamus, however, marker genes of multiple thalamic nuclei are enriched (*LHX6, CARTPT, TAC3*). About 36% of Subcluster 20 excitatory neurons are from the cortex and these are highly enriched for ASD genes (96.4% reaching nominal significance); most of them are identified as deep-layer corticothalamic, including L6b and L6 CT neurons (Supplementary Table 2), and they show high expression of marker genes including *BCL11B, FOXP2*, and *FEZF2*.

Subcluster 23 is also enriched for subcortical neurons, but with a higher fraction coming from thalamus (27.6% of total); the striatum is also overrepresented. The cells dissected from the thalamus were substantially enriched for ASD genes (67.3% reaching nominal significance). While subcluster 23 was enriched for general thalamic markers, including *SLC17A6* (VGLUT2), *SYNPR*, and *TAC1*, no individual thalamic nucleus was implicated by the marker genes. The cortical neurons within the cluster were less enriched for ASD genes (49.9% reaching nominal significance) and expressed markers of upper layers (L2/3: *SATB2, POU3F2*).

Subcluster 11 is composed of a mixture of neurons from the cortex (42.7% of cells; 57.0% reaching nominal significance for ASD enrichment), thalamus (20.1% of cells; 78.6% ASD enriched), and hypothalamus (19.7% of cells; 47.9% ASD enriched). The cortical neurons expressed markers of upper layers (L2/3: *SATB2, POU3F2*). No individual nucleus of the thalamus or hypothalamus was implicated by the marker genes and comparison to HypoMap showed enrichment across multiple hypothalamic nuclei with the addition of the Preoptic area over Subcluster 20.

## Discussion

Using a novel data integration approach, we assess ASD-associated gene enrichment across cell types, brain regions, and development in the 1^st^ and 2^nd^ trimester. We observe some clear results: there is strong enrichment in neurons of the second trimester, especially excitatory neurons. These findings are in keeping with prior analyses based on cortical data,[8, 23, 34, 35] limiting the scope of radically different results in other brain regions.

To understand whether specific brain regions are involved, we focused on excitatory neurons in the 2^nd^ trimester. Broadly, we see enrichment of ASD-associated genes across all brain regions (Fig. 3) and all excitatory neurons subtypes (Fig. 4). This could be interpreted as suggesting that ASD arises as a pan-neuronal phenomenon, i.e., a perturbation distributed across many neurons throughout the brain (e.g., an impairment at the level of synaptic plasticity). We cannot exclude this possibility; in fact, it would fit with the lack of focal lesions leading to ASD and diffuse findings from neuroimaging.[36, 37] However, this pan-neuronal enrichment may also represent limited regional and cell type resolution. In keeping with this possibility, genes associated with frontotemporal dementia and spinocerebellar ataxia are enriched in both prefrontal cortex and cerebellum, albeit with a greater degree of enrichment in the expected brain region.[38]

We see clear evidence of ASD-associated gene enrichment in the cortex, with three of the six ASD enriched clusters being cortical (clusters 1, 5, and 10). Across these, 95.9% of excitatory neurons are from the cortex compared with 75.7% across all excitatory neurons (1.3-fold enriched). The temporal cortex is the most enriched, accounting for 18.3% of excitatory neurons in the three cortical clusters compared to 7.1% across all excitatory neurons (2.6-fold enriched). Surprisingly, excitatory neurons from the prefrontal cortex (PFC) are underrepresented, making up 3.5% of the three clusters compared to 15.6% of all neurons (0.2-fold), however, of the PFC neurons that are in the three clusters, almost all of them are ASD enriched (92.5% in PFC; equivalent metric for temporal cortex is 69.5%).

Excitatory neurons from the hippocampus show the greatest degree of enrichment prior to correction for brain region (Fig. 3) and are highly enriched in cluster 5 (Fig. 4), where they account for 5.0% of excitatory neurons compared to 1.6% of all excitatory neurons (3.1-fold enriched). However, these data are derived from a single sample at GW18, so caution is warranted in taking this at face value. There has been substantial interest in investigating the role of the hippocampus in ASD in rodent models,[39, 40] especially in the context of synaptic plasticity and excitatory/inhibitory balance. In contrast, relatively little human-derived data exist to assess the role of the hippocampus.[36, 41]

We observe clear enrichment for cells in the 2^nd^ trimester thalamus (Fig. 2), driven by excitatory neurons (Fig. 3), which are enriched (2.5-fold enriched; 18.5% in cluster compared to 7.5% of all excitatory neurons) in all three of the subcortical ASD-enriched clusters (Fig. 4). We lack the resolution to distinguish specific thalamic nuclei,[42] however, cortical neurons in the same three clusters are enriched for markers of deep-layer corticothalamic neurons. We note that similar enrichment of ASD-associated genes is observed in the spatial transcriptomic analysis of the prenatal human brain[43] and adult mouse brain (Yu *et al*. 2026). The thalamus has received relatively limited attention in ASD research,[44] however, evidence of thalamocortical hyperconnectivity was described from resting-state functional MRI of 81 autistic individuals compared to 71 age- and sex-matched controls.[45] Supportive evidence for thalamocortical hyperconnectivity comes from organoid models of the ASD-associated 22q11.2 microdeletion.[46] Structural MRI data from 373 male autistic individuals compared to 384 male controls showed expanded surface area of the pulvinar nucleus in the thalamus.[47] Homozygous knockout of *CNTNAP2* has been implicated in ASD;[48] analysis of equivalent *Cntnap2*^*-/-*^ mice identified hyperactivity of reticular thalamic nucleus.[49] In contrast, the homozygote mouse model (*Shank3*^-/-^) of human *SHANK3* haploinsufficiency was found to have hypoactivity of reticular thalamic nucleus[50] and male Fragile-X mice (*Fmr1*^*-*^) had impaired thalamic burst firing in the lateral geniculate nucleus.[51]

We also saw evidence of enrichment for excitatory neurons of the hypothalamus in the three subcortical clusters (3.6-fold enriched; 18.2% in cluster compared to 4.8% of all excitatory neurons; Fig. 4). Like the thalamus, the hypothalamus has not been a prominent target of ASD research;[52] however, it contains sex-differential neuronal populations that drive sex-differential behaviours, particularly in nuclei such as the ventrolateral subdivision of the ventromedial hypothalamus (VMHvl).[53]

To enable these regional insights, we required a statistically robust method to quantify the regional variances and reveal the biological signal contained in the region labels themselves, as well as after accounting for the regional variances. To achieve this, we developed STARMAPS, which uses the idea of task-specific subspace perturbation, which has been utilized in the language/computer vision applications. This model provides a natural way to check the amount of perturbation, and the relationship between latent spaces.

We note limitations of this work. Our assessment of ASD-associated gene enrichment relies on the underlying assumption that higher expression predicts a causal role. This makes intuitive sense at the extremes (e.g., a cell is less likely to mediate ASD if the identified genes are not expressed) but is not a foregone conclusion. We also assume some degree of convergence across genes, rather than assessing each gene independently; this is supported by analyses of gene function.[8] Most of our results are based on thirteen individual brains in the 2^nd^ trimester data, with smaller subsets in specific regions; while we have used STARMAPS to minimize batch effects and highlighted where results are limited to very few samples, this sample size is limiting. We also lack samples during the 3^rd^ trimester or postnatal development or from the cerebellum, giving and incomplete map of spatiotemporal enrichment. The STARMAPS model could also be improved by utilizing prior knowledge about the correlation between regions to improve the accuracy of task-specific perturbation estimation.

## Conclusion

Recently, large-scale scRNA-seq dataset of human fetal brain with multiple regional samples enables exploration of areal signatures of gene expression. Previous results have shown that, while cell type is the major driving force of segregation, region labels also contribute significantly to the scRNA expression profiles.[54] It has also been shown that both developmental stage and cell type contribute to the variance of ASD risk gene expression patterns.[3] We replicate prior findings implicating excitatory neurons in the 2^nd^ trimester in ASD and extend these to implicate the cortex, thalamus, and hypothalamus.

## Acknowledgements

We are grateful to the families who donated the initial samples in the Bhaduri and Eze papers on which these analyses are performed. This work was supported by a grant from the Simons Foundation Autism Research Initiative Sex Differences Collaboration (SFARI #736613 to S.J.S.), the National Institute of Mental Health (grant numbers R01MH125516 to S.J.S, R01MH129751 to S.J.S., R01MH129725 to K.R., and R01MH123184 to K.R.), HDRUK: Health Data Research UK QQ2 Molecules to Health Records Driver Programme to S.J.S, NARSAD Young Investigator grants (31904) to X.W. and Seaver Foundation Fellow to X.W.

## Materials and Methods

The advance of scRNA-sequencing technique enables fine-grained sequencing results, precise up to the brain region in human fetal brains. This raises the question about the regional effect of scRNA gene expression; however, most current works are conducted in a heuristic manner.[55–58] In this paper, our goal is to improve upon existing approaches in order to quantify the regional variances and reveal the biological signal contained in the region labels themselves, as well as after accounting for the regional variances in a statistically solid/robust framework. The problem is of similar flavor to the batch effect correction, or dataset integration, such as cFIT,[22] but the task is more subtle: the regional effect is not a gene-wise shift (which is a common assumption for the batch/dataset effect) but rather a global and simultaneous perturbation on a large number of genes. Due to the abundance of regions, it is more prone to overfitting. Thus, compared to the dataset integration task, the interpretability of the model is of vital importance, as is the need to account for the batch and regional effect at the same time.

As such, inspired by cFIT framework, we developed a new method, STARMAPS (Sparse Task-specific Analysis for Revealing Molecular Associations in Particular Single-cell datasets) with more flexibility and interpretability. We use the idea of task-specific subspace perturbation, which has been utilized in language/computer vision applications. Building upon the non-negative matrix factorization framework, we assume that each region is associated with a latent factor space, and then we regularize directly on the (approximate) subspace distance between those factor spaces. STARMAPS is flexible with more freedom on parameters, while we can control the task-specific perturbation easily by changing the regularization parameters. This model also provides a natural way to check the amount of perturbation, and the relationship between latent spaces.

### STARMAPS model for data-specific effects

With the non-negative matrix factorization framework, we add a sparse perturbation on different regions. Suppose we know the region labels and batch labels, the expression matrix associated with batch *j* and region *r* is 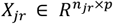. The objective function is:

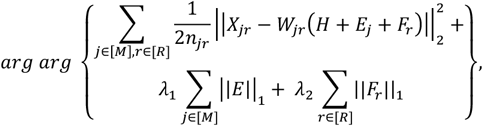

subject to

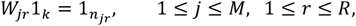

where *H, E*_*j*_, *F*_*r*_ ∈*R*^*k*×*p*^ and 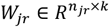, with *k* being the predetermined number of hidden dimensions.

Here, we assume the subspace associated with each (batch, region) can be decomposed into three additive parts: the shared subspace across all data, the sparse batch-specific perturbation, and the sparse region-specific perturbation. The *L*_*1*_ penalty terms are added to control the size of the perturbation matrices. We add the constraints that *H* and all *W*_*jr*_s and *H* + *E*_*j*_ + *F*_*r*_s are non-negative matrices.

Details of the algorithm are listed below.

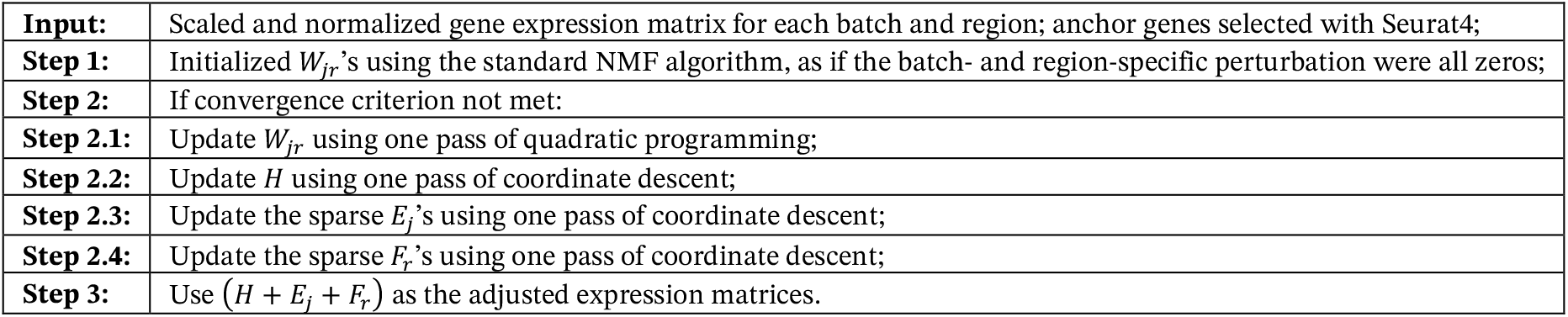

### Removal of the regional effect

The first step is to validate the success of removing the regional effect of the proposed algorithm. Intuitively, if the regional variances were removed indeed, conducting clustering on the reconstructed membership matrices would give a more balanced coupling between region labels and cluster labels. Based on such reasoning, we introduce the following evaluation criterion. Suppose some clustering algorithm divides all the n cells into *K* clusters, labeled by *c*_*i*_ ∈{1, …, *K*}, 1 ≤ *i* ≤ *n*. For each region *r* ∈[*R*] and cluster *k* ∈[*K*], we calculate the proportion of the region *r* cells in cluster *k* among all the cluster *k* cells, and then measure the weighted total variance distance as

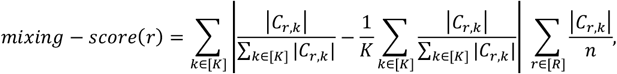

Where the cell group *C*_*r,k*_ is defined as

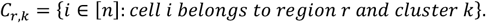

Based on the above rationale, a lower mixing score indicates removal of regional variance. In real implementation, we use a tree-based hierarchical clustering algorithm called MRTree.[22] We reported the mixing scores for each region, and with model regularization *λ* = 0, 10^-2^ and 10^-3^ as a bar plot shown in Fig. S1A. From this figure, we can see that smaller lambda (or less regularization on the regional perturbation) does remove the regional correlations in the membership matrices, building the ground of downstream reasoning.

#### Biological signals

Another important criterion to check is that STARMAPS does not eliminate the true biological signals in the reconstructed scRNA expression profiles. To see this, we use the three modelling options introduced above and conduct cross-validated classification analysis on the cell type labels, one of the most important cellular signatures, and of which we have the ground-truth labels from the original dataset. Precisely, in each run, we randomly select 1000 cells in the whole dataset as query data, and another non-overlapping 4000 cells as reference data. Then we use one shared-neighbour-based mapping algorithm[21, 59] to label the query dataset and compare it to the truth. Repeat for 10 times for each model setting, and report the average classification accuracy, as well as the half standard deviation. The results are shown in Fig. S1B. It is clearly seen that adding the regularization terms reduces the noises in learning true biological signals.

#### Amount of region and batch perturbations

For further illustration, we calculate the amount of region/batch perturbations, using an approximate measure of distance between linear subspaces

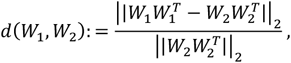

where the denominator is for normalization purposes. The above measure is widely used in the statistical literature and is shown to be equivalent to the cosine distance and the rotation distance between linear subspaces, as shown in.[60, 61] The denominator serves as a shared normalization factor. We plot the resulting subspace distance measures in Fig. S1C and S1D. We can see that the cortex areas go through the least amount of correction; in other words, the learned subspace perturbation structure can be approximately perceived as the divergence from the cortex subspace. Also note that compared to the regional perturbations, the batch effects are smaller in scale. (Notice that the only one large batch perturbation term comes from the age GW22, the age with the least sample size; as such, this perturbation term is likely to be noisier. The other batch perturbations are almost negligible, compared to the regional ones.)

### Prenatal brain datasets

We use two prenatal brain datasets in this paper, namely, Eze *et al*. 2021 and Bhaduri *et al*. 2021 single cell atlases.[20, 21] The Bhaduri atlas sequenced single-cell transcriptomes from micro-dissected regions of developing human brain tissue during the second trimester, which encompasses peak stages of neurogenesis. It contains cells from 17 forebrain, midbrain, and hindbrain regions from 13 individuals. In particular, the cortex region cells are labeled with more fine-grained labels: PFC, motor, somatosensory, parietal, temporal, and primary visual (V1) cortex. There are 698,820 cells in total, and we use a random subset of 100,000 in our analysis. The Eze atlas contains cells from 10 individuals during the first trimester of human development, spanning Carnegie stages (CS) 12 to 22, corresponding to gestational weeks 6–10, with a total sample size of 289,000. We use 50,000 of them in our analysis. We make the note that the Eze cells are at a substantial earlier developmental stage than the whole Bhaduri dataset.

### ASD risk genes

We use the results from Fu *et al*. 2022 to identify the ASD-associated genes.[3] These gene association results are based on joint statistical analysis of rare protein-truncating variants (PTVs), damaging missense variants, and copy number variants (CNVs) derived from exome sequencing of 63,237 individuals from ASD cohorts. We use the 185 genes at FDR <0.05 as the ASD risk genes for analysis in the rest of the paper.

### Enrichment Analysis with Odds Ratio

With ASD risk genes provided above, we calculate the ASD enrichment by odds ratio (OR) and use the associated p-values for its statistical significance. Before presenting more numerical details, we first describe the calculation of OR and its significance level in our setting.

Calculation of the ES using the two-by-two frequency table:

|  | # genes expressed | # genes not expressed |
| --- | --- | --- |
| ASD risk genes | $n_{11}$ | $n_{12}$ |
| Non-ASD risk genes | $n_{21}$ | $n_{22}$ |

The first step of the enrichment analysis is the calculation of the two-by-two frequency table, as illustrated in Table 1, of the conditions ASD risk genes/non-ASD risk genes, versus number of genes expressed/not expressed. The OR is then essentially the results of Fisher exact test of this contingency table fisher1922interpretation

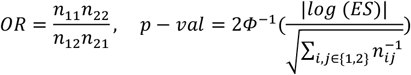

To complete the enrichment analysis, we need to clarify the criterion for “expressed” in Table 1. Use *X* to denote the original count dataset for one of the cell developmental stages, 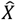 to denote the corresponding reconstructed expression profile, and *x*_*i*_ to denote the reconstructed profile of cell *i*. We identify one gene *g* as “expressed” in a cell subgroup *S*, when *g* is expressed in more than 25% of the cells in *S*. We identify *g* as expressed in a cell, when:

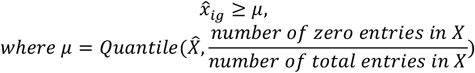

Here the choice of *μ* acts as the threshold for “expression”, such that the reconstructed dataset shares the same number of “expressed” entries as the original dataset. One such thresholding step in expression (2) is necessary since the reconstructed dataset is no longer sparse.

Both Bhaduri and Eze dataset come with pre-assigned cell labels, and we use these labels to identify various subgroups in each region. Following such subgroup construction, we proceed to calculating the ES and associated p-values of all the subgroups and identify the “enriched sub-groups” by OR > 1 and p < 0.05. Then, for each region, we calculate the percentage of cells belonging to the “enriched sub-groups”. Note that here we calculate the percentage of cells instead of groups, since the group sizes are highly heterogeneous, in which case percentage of cells reflect the true extent of enrichment in a more accurate and stable manner.

First, we demonstrated that, with proper choices of regularization parameters (Fig. S1), the proposed algorithm removes the regional variances in the membership matrix *W*_*jr*_, while preserving the true biological signals. We also evaluate the amount of regional variance for all the regions.

Second, we use the MRtree algorithm, a tree-based hierarchical clustering algorithm introduced in Peng *et al*. 2021,[22] to identify cell subgroups with proper resolution, in a robust manner.

Third, we run odds ratio enrichment on the identified subgroups, and report the biological markers (odds ratio, marker gene, regional compositions, among others) on these subgroups, and discuss the potential significance in ASD research.

As discussed in the Results, most signals lie in the excitatory neurons (and, to a lesser extent, inhibitory neurons). As such, we use the excitatory neurons of Bhaduri data in the following analysis.

### Cell type hierarchy reconstruction via MRTree

Here, we obtain a low-dimensional embedding of the membership matrix, and then apply the MRTree method proposed by Peng *et al*.[22] to identify subgroups in the regional-corrected expression profiles.

Results in the previous section suggest us to use the *λ* = 10^-3^ model, to achieve the desired goals of removing regional effects and preserving biological signals; the preceding analysis will be based on this parameter choice. In Figure 3, we plot the UMAP embedding of the excitatory neurons, colored by subtypes, to give a visualization of the cell membership matrix.

### ASD gene enrichment analysis for the identified subgroups

Finally, we conduct the ASD gene enrichment on the identified subgroups. The clusters with p-val < 0.05 are indicated in Figure 4, and the enriched clusters are also marked with ∗∗ for p-val < 0.01 and ∗ for p-val < 0.05. The summarized p-values and the multiple-testing adjusted p-values are in Supplementary Table 3. The enriched cell subgroups are highlighted on the UMAP visualization in Figure 4A and summarize their regional composition in Figure 4B and C, in both the region labels and the major region labels. The marker genes and the expression heatmap of the enriched clusters are in Supplementary Table 2.

### Mapping subgroups to a hypothalamic dataset

We mapped subclusters to a unified single cell dataset of the mouse hypothalamus, HypoMap, using the Seurat label transfer pipeline (Fig. S5). First, we found transfer anchors between our dataset and HypoMap using the “SCT” normalization method, with HypoMap downsampled by 1:100 to fit within memory constraints. We next employed these anchors to transfer our dataset’s subcluster labels to the hypothalamic cells, generating a prediction score of each subcluster label for each cell. We aggregated these prediction scores by mean across cells within each hypothalamic nucleus label to show the degree of match between each of our subclusters and the hypothalamic nuclei, plotted as a heatmap using the heatmaply package.

