## Supplemental Figures and Tables for "Spatiotemporal analysis of autism gene enrichment implicates cortex, thalamus, and hypothalamus"

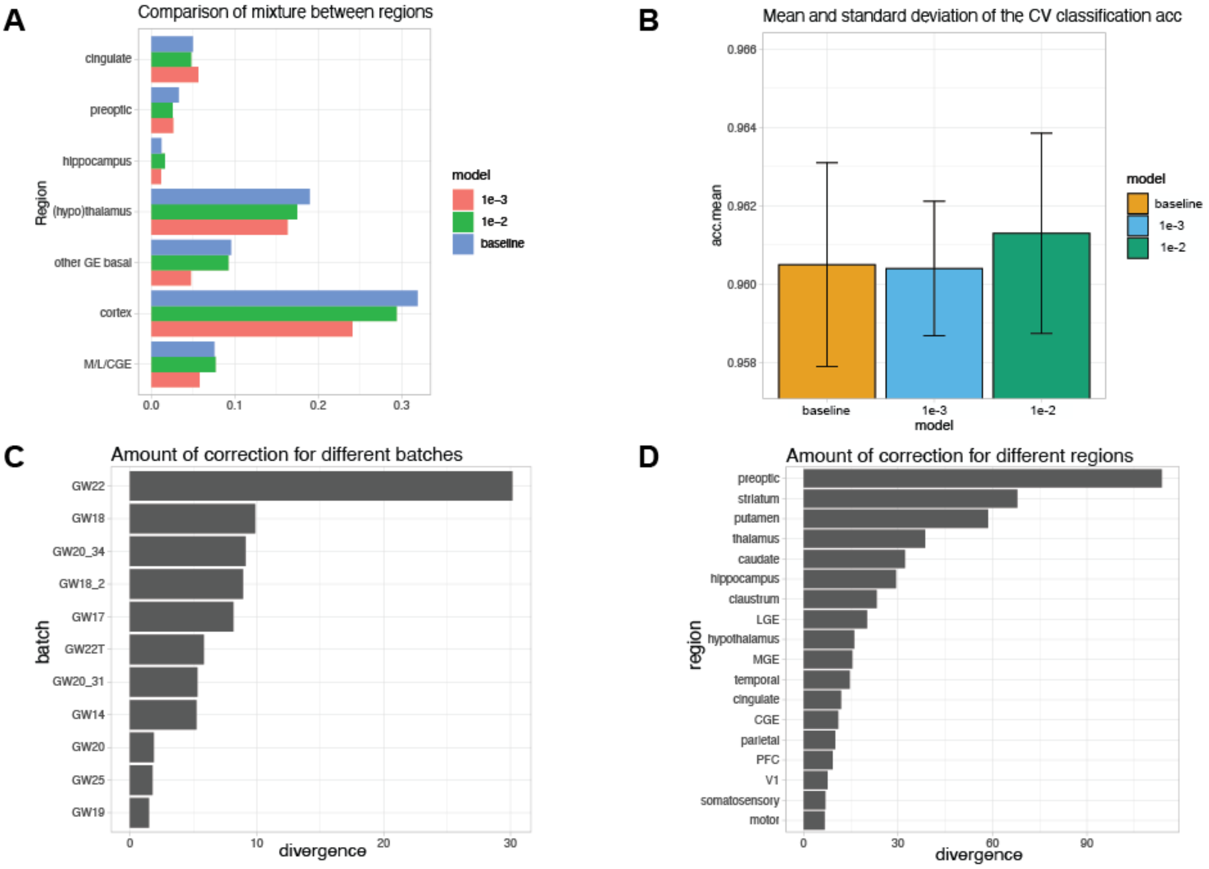


**Supplemental Figure S1:** Parameter selection and correction of sample and regional effects. **A)** Mixing scores for each area with model regularization $\lambda = 0, 10^{-2}$ and $10^{-3}$; smaller lambda (or less regularization on the regional perturbation) does remove the regional correlations in the membership matrices, building the basis of downstream reasoning. **B)** Mean and the standard deviation of the CV classification ACC. **C)** Amount of correction for different batches, $\lambda = 10^{-3}$. **D)** Amount of correction for different regions, $\lambda = 10^{-3}$.

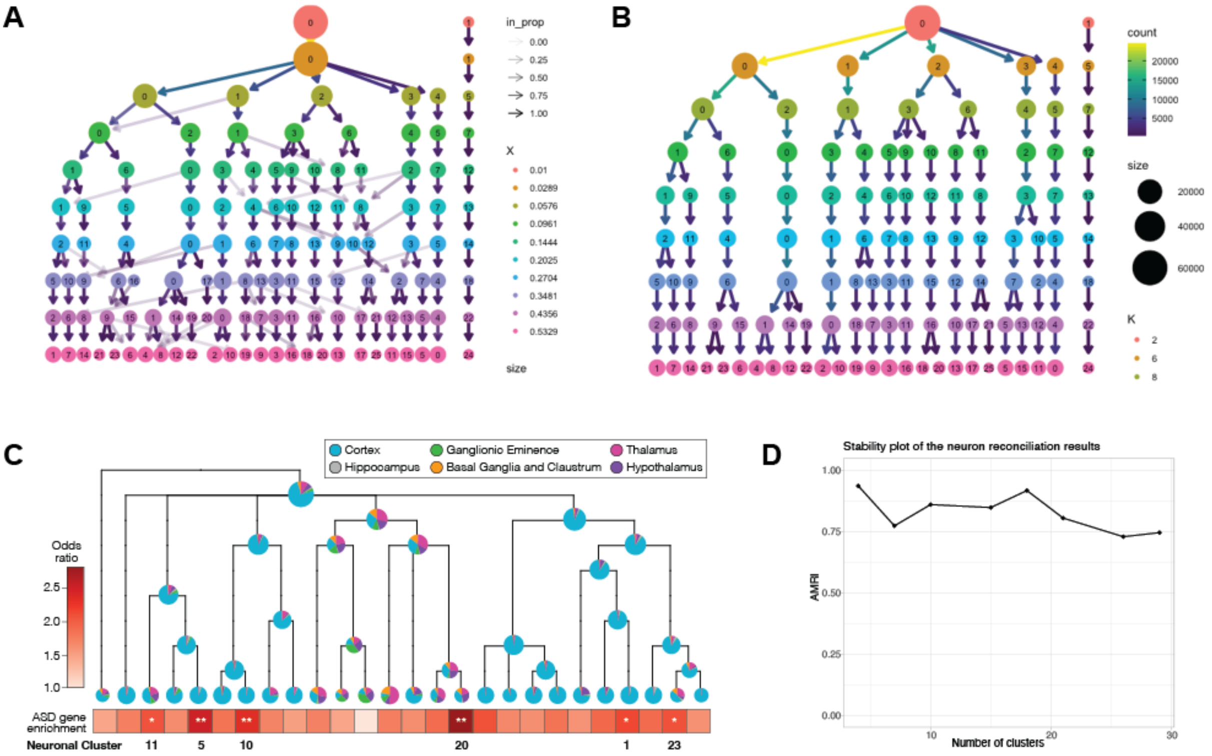


**Supplemental Figure S2.** MRTree clustering of excitatory neurons. **A)** The clustering results from FindMarker() (Seurat v5.0.3), estimated separately for each layer. **B)** The reconstructed hierarchical tree using MRTree. **C)** Regional composition of the nodes on each level along with ASD risk gene enrichment of the 26 leaf nodes. **D)** Stability plot of the number of clusters, indicating that the clustering results are robust to the parameter choices. The leaf level of the MRTree produces 26 clusters for the excitatory neurons, which was used for subsequent analyses.

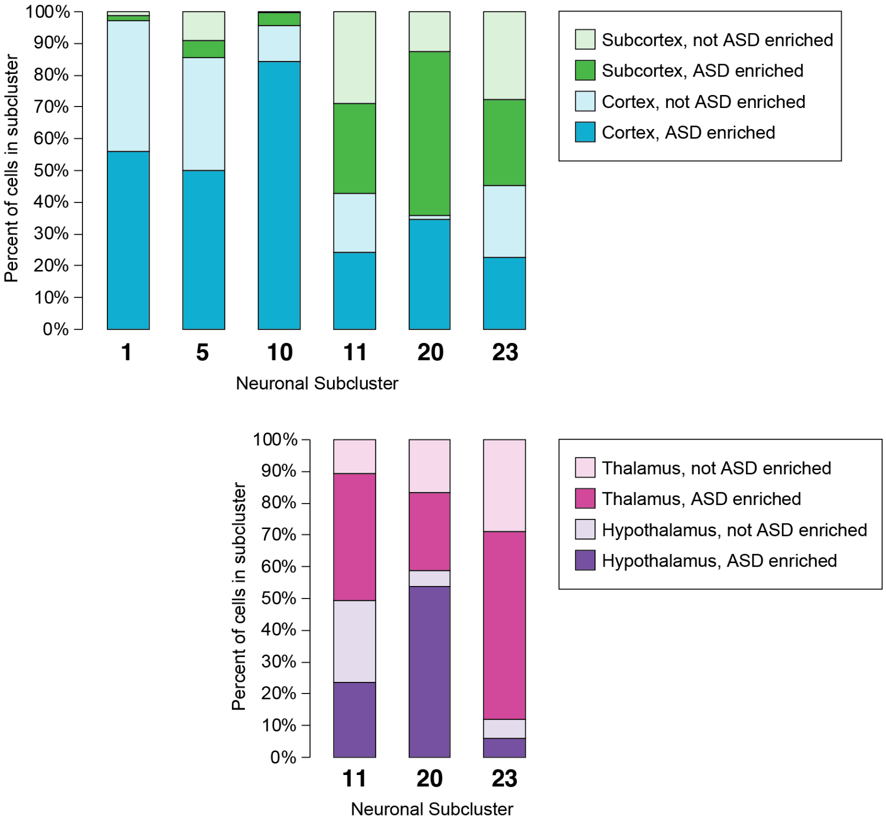


**Supplemental Figure S3.** Enrichment of ASD-associated genes. Excitatory neurons in the cortex (blue) and subcortex (green) are colored by the presence (darker shades) or absence (lighter shades) of enrichment for ASD-associated genes. Similarly, excitatory neurons in the thalamus (pink) and hypothalamus (purple) are colored by the presence (darker shades) or absence (lighter shades) of enrichment for ASD-associated genes.

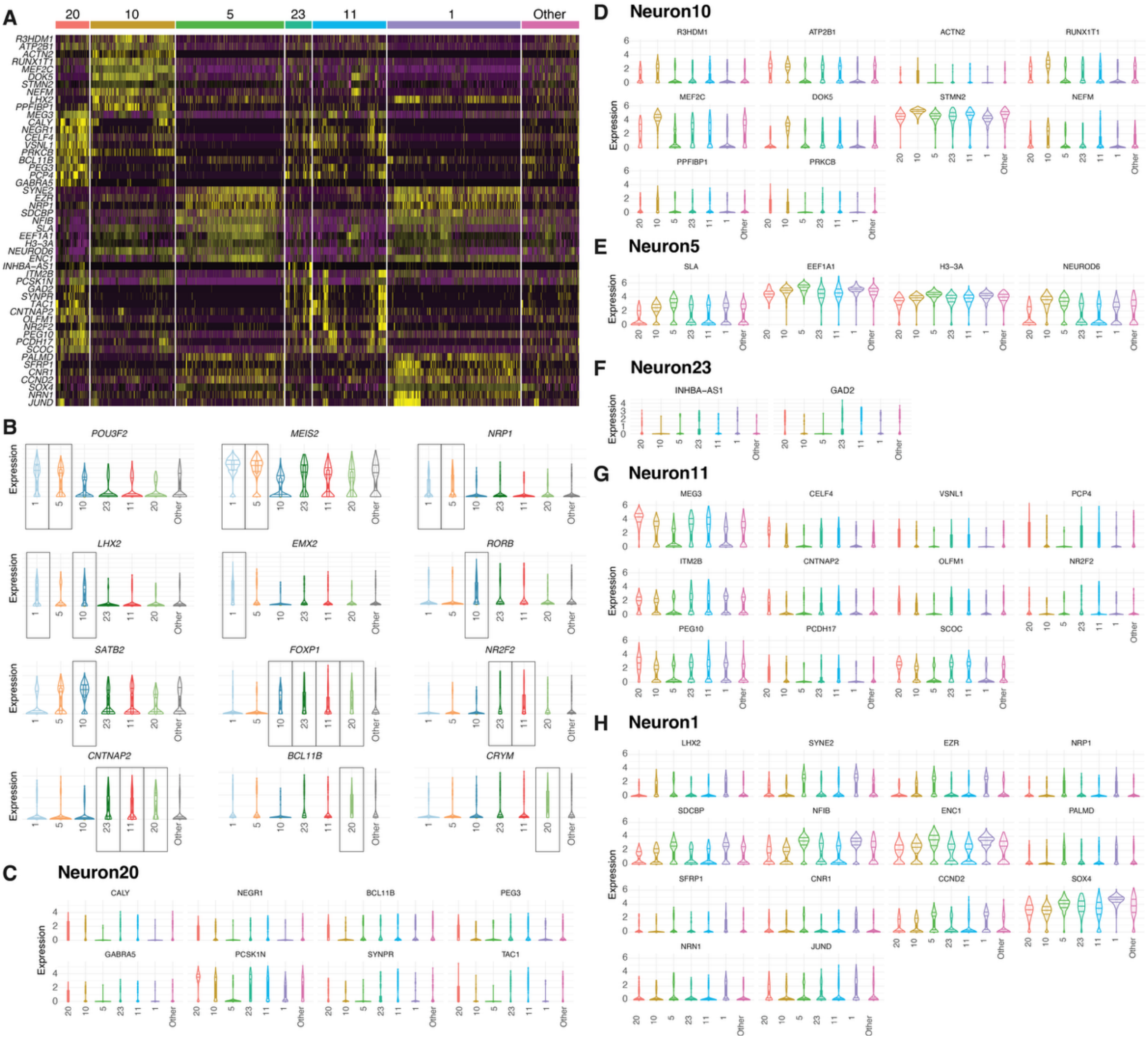


**Supplementary Figure S4. Marker genes for ASD-enriched neuron subtypes. A.** Heatmap of subtype specific marker gene expression for ASD-enriched neuron subtypes. **B**. Violin plot of known cell type markers and their expression in the ASD-enriched neuron subtypes. **C-H.** Violin plot of subtype specific marker gene expressions for the ASD-enriched neuron subtypes.

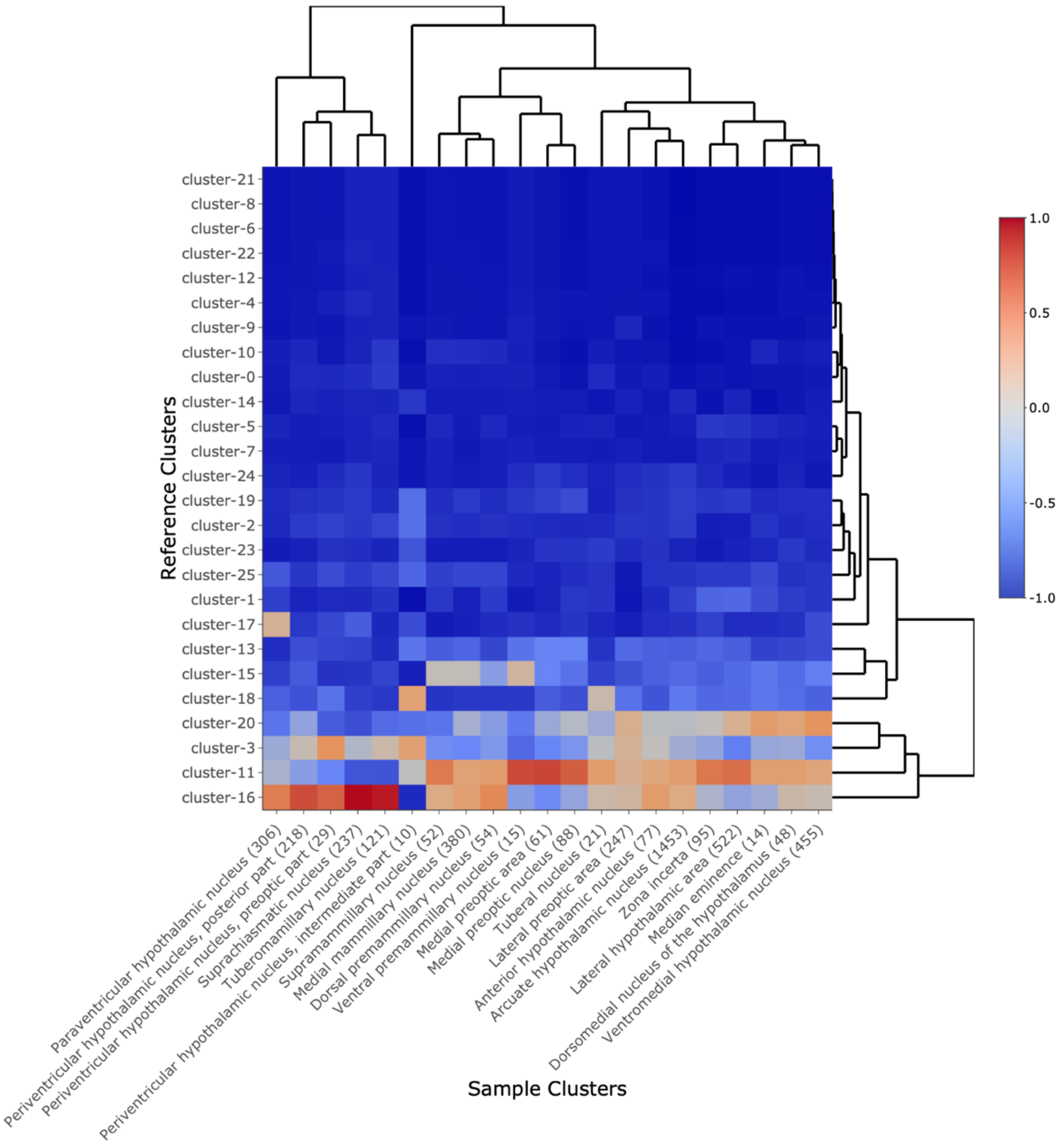


1,2

**Supplemental Figure S5. Mapping of subclusters to hypothalamic clusters from HypoMap.** Developing human excitatory neuronal subclusters were projected onto hypothalamic clusters in HypoMap, with prediction scores for cluster matches shown as a heatmap, scaled by hypothalamic cluster. Subcluster 20, which showed high enrichment for ASD-associated genes and a large proportion of neurons from the hypothalamus, matched strongly with several regions including the median eminence, dorsomedial nucleus of the hypothalamus, and ventromedial hypothalamic nucleus. Numbers of cells in each hypothalamic cluster are indicated in parentheses.

| Major Regions | Regions |
|---|---|
| Cortex | PFC |
| | Motor |
| | Somatosensory |
| | Parietal |
| | Temporal |
| | V1 |
| | Cingulate |
| Hypothalamus | Preoptic |
| | Hypothalamus |
| Basal Ganglia and Claustrum | Striatum |
| | Putamen |
| | Caudate |
| | Claustrum |
| Ganglionic Eminence | LGE |
| | CGE |
| | MGE |
| Thalamus | Thalamus |
| Hippocampus | Hippocampus |

**Supplemental Table 1.** Major regions and corresponding regions in the Bhaduri dataset.

| Gene | p-value | Avg_log2FC | Adj p-value | cluster | gene.name |
|---|---|---|---|---|---|
| **ENSG000000489912** | 0.000000e+00 | 1.1413112 | 0.000000e+00 | 10 | *R3HDM1* |
| **ENSG000000709612** | 0.000000e+00 | 0.8624473 | 0.000000e+00 | 10 | *ATP2B1* |
| **ENSG00000077522** | 0.000000e+00 | 1.3305071 | 0.000000e+00 | 10 | *ACTN2* |
| **ENSG000000791022** | 0.000000e+00 | 1.5612273 | 0.000000e+00 | 10 | *RUNX1T1* |
| **ENSG000000811893** | 0.000000e+00 | 1.6275406 | 0.000000e+00 | 10 | *MEF2C* |
| **ENSG000001011343** | 0.000000e+00 | 1.6136567 | 0.000000e+00 | 10 | *DOK5* |
| **ENSG000001044354** | 0.000000e+00 | 0.6438235 | 0.000000e+00 | 10 | *STMN2* |
| **ENSG000001047222** | 0.000000e+00 | 1.3676967 | 0.000000e+00 | 10 | *NEFM* |
| **ENSG00000106689** | 2.027951e-290 | 1.1310808 | 6.923425e-287 | 1 | *LHX2* |
| **ENSG000001108411** | 0.000000e+00 | 1.4003115 | 0.000000e+00 | 10 | *PPFIBP1* |
| **ENSG000002145482** | 3.230490e-109 | 0.9195507 | 1.102889e-105 | 11 | *MEG3* |
| **ENSG000001306431** | 1.085761e-296 | 2.1174005 | 3.706786e-293 | 20 | *CALY* |
| **ENSG000001722602** | 1.435167e-285 | 1.4875034 | 4.899660e-282 | 20 | *NEGR1* |
| **ENSG000001014892** | 2.514454e-106 | 0.9071810 | 8.584347e-103 | 11 | *CELF4* |
| **ENSG000001630321** | 4.197787e-87 | 0.9390323 | 1.433125e-83 | 11 | *VSNL1* |
| **ENSG00000166501** | 1.474478e-232 | 0.9722231 | 5.033867e-229 | 10 | *PRKCB* |
| **ENSG000001271523** | 2.362602e-257 | 0.8032720 | 8.065924e-254 | 20 | *BCL11B* |
| **ENSG000001983002** | 4.402825e-255 | 1.6076137 | 1.503125e-251 | 20 | *PEG3* |
| **ENSG000001830361** | 4.997348e-85 | 0.7456092 | 1.706095e-81 | 11 | *PCP4* |
| **ENSG00000186297** | 4.564354e-250 | 1.2517788 | 1.558271e-246 | 20 | *GABRA5* |
| **ENSG00000054654** | 0.000000e+00 | 1.6285767 | 0.000000e+00 | 1 | *SYNE2* |
| **ENSG00000092820** | 0.000000e+00 | 1.3823014 | 0.000000e+00 | 1 | *EZR* |
| **ENSG00000099250** | 0.000000e+00 | 1.3671946 | 0.000000e+00 | 1 | *NRP1* |
| **ENSG00000137575** | 0.000000e+00 | 1.0601902 | 0.000000e+00 | 1 | *SDCBP* |
| **ENSG000001478621** | 0.000000e+00 | 1.0967165 | 0.000000e+00 | 1 | *NFIB* |
| **ENSG00000155926** | 0.000000e+00 | 1.3214578 | 0.000000e+00 | 5 | *SLA* |
| **ENSG00000156508** | 0.000000e+00 | 0.6551033 | 0.000000e+00 | 5 | *EEF1A1* |
| **ENSG00000163041** | 0.000000e+00 | 0.5559999 | 0.000000e+00 | 5 | *H3-3A* |
| **ENSG000001646002** | 0.000000e+00 | 0.7928197 | 0.000000e+00 | 5 | *NEUROD6* |
| **ENSG00000171617** | 0.000000e+00 | 1.0974793 | 0.000000e+00 | 1 | *ENC1* |
| **ENSG00000224116** | 4.439962e-185 | 1.4395486 | 1.515803e-181 | 23 | *INHBA-AS1* |
| **ENSG00000136156** | 3.066431e-153 | 0.9970797 | 1.046880e-149 | 11 | *ITM2B* |
| **ENSG000001021096** | 3.532679e-97 | 0.9080235 | 1.206057e-93 | 20 | *PCSK1N* |
| **ENSG000001367501** | 7.764232e-51 | 1.2086886 | 2.650709e-47 | 23 | *GAD2* |
| **ENSG000001636302** | 2.397409e-86 | 0.4174026 | 8.184753e-83 | 20 | *SYNPR* |
| **ENSG000000061282** | 1.368571e-101 | 1.6067870 | 4.672301e-98 | 20 | *TAC1* |
| **ENSG000001744691** | 9.766431e-82 | 0.9290922 | 3.334259e-78 | 11 | *CNTNAP2* |
| **ENSG00000130558** | 2.834704e-29 | 0.4641852 | 9.677680e-26 | 11 | *OLFM1* |
| **ENSG00000185551** | 5.305655e-304 | 2.1514963 | 1.811351e-300 | 11 | *NR2F2* |
| **ENSG000002422651** | 1.563009e-101 | 1.2027769 | 5.336111e-98 | 11 | *PEG10* |
| **ENSG000001189461** | 1.606920e-85 | 0.9130276 | 5.486026e-82 | 11 | *PCDH17* |
| **ENSG000001531302** | 1.717792e-84 | 0.6648287 | 5.864541e-81 | 11 | *SCOC* |
| **ENSG00000099260** | 0.000000e+00 | 1.2670068 | 0.000000e+00 | 1 | *PALMD* |
| **ENSG00000104332** | 0.000000e+00 | 1.4003975 | 0.000000e+00 | 1 | *SFRP1* |
| **ENSG00000118432** | 0.000000e+00 | 1.2278695 | 0.000000e+00 | 1 | *CNR1* |
| **ENSG00000118971** | 0.000000e+00 | 1.0545852 | 0.000000e+00 | 1 | *CCND2* |
| **ENSG000001247661** | 0.000000e+00 | 0.8178123 | 0.000000e+00 | 1 | *SOX4* |
| **ENSG00000124785** | 0.000000e+00 | 1.5987086 | 0.000000e+00 | 1 | *NRN1* |
| **ENSG00000130522** | 0.000000e+00 | 1.4097674 | 0.000000e+00 | 1 | *JUND* |

**Supplementary Table 2.** Marker genes of ASD enriched neuron subtypes.

| Cluster | Odds Ratio | p-value | Adj p-value |
|---|---|---|---|
| **Neuron10** | 2.187 | 0 | 0 |
| **Neuron20** | 2.808 | 0 | 0 |
| **Neuron5** | 2.405 | 0 | 0.002 |
| **Neuron23** | 1.822 | 0.001 | 0.023 |
| **Neuron11** | 1.801 | 0.002 | 0.035 |
| **Neuron1** | 1.914 | 0.002 | 0.035 |
| Neuron12 | 1.836 | 0.006 | 0.107 |
| Neuron18 | 1.560 | 0.008 | 0.145 |
| Neuron7 | 1.605 | 0.021 | 0.353 |
| Neuron2 | 1.492 | 0.029 | 0.462 |
| Neuron6 | 1.576 | 0.033 | 0.493 |
| Neuron8 | 1.490 | 0.055 | 0.769 |
| Neuron15 | 1.229 | 0.396 | 1.000 |
| Neuron3 | 1.384 | 0.177 | 1.000 |
| Neuron19 | 1.313 | 0.195 | 1.000 |
| Neuron13 | 1.343 | 0.129 | 1.000 |
| Neuron16 | 1.216 | 0.326 | 1.000 |
| Neuron14 | 1.144 | 0.514 | 1.000 |
| Neuron0 | 1.363 | 0.144 | 1.000 |
| Neuron4 | 1.196 | 0.378 | 1.000 |
| Neuron9 | 1.034 | 0.836 | 1.000 |
| Neuron22 | 1.281 | 0.233 | 1.000 |
| Neuron21 | 1.174 | 0.407 | 1.000 |
| Neuron17 | 0.347 | 0.000 | 0.000 |
| Neuron24 | 0.970 | 1.000 | 1.000 |
| Neuron25 | 0.966 | 1.000 | 1.000 |

**Supplementary Table 3**. Enrichment of excitatory neuron subtypes, including odds ratios, p-values, and adjusted p-values. The top six excitatory neuron subtypes have an odds ratio greater than 1 and an adjusted p-value less than 0.05.
